# A constitutively expressed antifungal peptide protects *Tenebrio molitor* during a natural infection by the entomopathogenic fungus *Beauveria bassiana*

**DOI:** 10.1101/284778

**Authors:** Sevasti Maistrou, Véronique Paris, Annette B. Jensen, Jens Rolff, Nicolai V. Meyling, Caroline Zanchi

**Author notes:** Westfälische Wilhelms Universität, Institute for Evolution & Biodiversity, Hüfferstraße 1, 48149 Münster, Germany.

## Abstract

Antimicrobial peptides have been well studied in the context of bacterial infections. Antifungal peptides have received comparatively less attention. Fungal pathogens of insects and their hosts represent a unique opportunity to study host-pathogen interactions due to the million of years of co-evolution they share. In this study, we investigated role of a constitutively expressed thaumatin-like peptide with antifungal activity expressed by the mealworm beetle Tenebrio molitor, named Tenecin 3, during a natural infection with the entomopathogenic fungus Beauveria bassiana. We monitored the effect of the expression of Tenecin 3 on the survival of infected hosts as well as on the progression of the fungal infection inside the host. Finally, we tested the activity of Tenecin 3 against B. bassiana. These findings could help improving biocontrol strategies and help understanding the evolution of antifungal peptides as a defense mechanism.

## 1. Introduction

Fungal pathogens infect a great variety of host species including humans, such as *Candida albicans, Naegleria fowleri, Aspergillus* sp., or are pests of economically important crops, such as *Fusarium oxysporum* (Fisher et al., 2012). Fungal pathogens of insects, called entomopathogenic fungi, share millions of years of coevolution with their insect hosts (Boomsma et al., 2014; Joop & Vilcinskas, 2016). These fungi play an important role in controlling insect populations in natural ecosystems by causing epizootics (Hesketh et al., 2010), thus exerting a strong selection pressure on their hosts. The relatively short life-cycles of most insects and entomopathogenic fungi, as well as the knowledge of virulence factors of the fungus (Samuels et al., 1988; Hajek & St. Leger, 1994; Valero-Jimenez et al., 2016) and immune reactions of the insect host (Butt et al., 2016), make these organisms a attractive models for the study of host-pathogen interactions. Because most entomopathogenic fungi are necrotrophic parasites (i.e. need to kill their host in order to complete their life cycle), they have been extensively studied in the context of biocontrol (Lacey et al., 2015).

The entomopathogenic fungus *Beauveria bassiana* (Ascomycota: Hypocreales) infects a broad range of hosts (Ferron, 1978; Inglis et al., 2001) such as ants (Broome et al., 1976), termites (Culliney & Grace, 2000), agricultural pests such as grasshoppers (Bidochka & Khachatourians, 1991) and arthropod vectors of human diseases (Clark et al., 1968; Kirkland et al., 2004; Marti et al., 2005). Once *B. bassiana* conidia land on a potential host, they attach to the cuticle, germinate and penetrate the cuticular barrier by a combination of mechanical pressure and enzymatic secretions (Ortiz-Urquiza & Keyhani, 2013). When the fungus reaches the hemolymph, it differentiates into yeast-like, thin-walled blastospores or hyphal bodies, which divide and exploit host nutrients. After the death of the host and after catabolizing its last nutrients, hyphae are produced which breach the cuticle from the inside and form conidia at the surface of the cadaver, thereby completing the asexual life cycle of the fungus (Ortiz-Urquiza and Keyhani 2013; Pedrini et al. 2013).

Once conidia come in contact with the host, the first line of defense is the cuticle that forms a mechanical barrier which, in combination with defenses such as cuticular hydrocarbons (Boyle & Cutler, 2012; Lopes et al., 2015), and protease inhibitors (Li et al., 2012), have been well studied for their role in host resistance (Ortiz-Urquiza & Keyhani, 2013). When the cuticle has been breached, the fungus will face the effectors of the insect immune system, where immune reactions relying on immune cells and the phenoloxidase pathway, such as phagocytosis (Gillespie et al., 2000), encapsulation and nodule formation (Lord et al., 2002), have been shown to play a central role in the suppression and clearance of *B. bassiana* from the host hemolymph.

However, insects also possess a battery of antimicrobial peptides (AMPs), small cationic peptides showing a relatively broad spectrum of antimicrobial activity ranging from Gram positive and/or negative bacteria, protozoans, viruses, and fungi (Bulet et al., 1999; Hancock, 2000; Peschel & Sahl, 2006; Mylonakis et al., 2016).

AMPs have attracted a lot of attention in the context of bacterial infections (Yokoi et al., 2012; Johnston et al., 2014) and because of their potential as new therapeutical agents (Zasloff, 2002). Some AMPs have shown antifungal activity either while being also antibacterial (Levashina et al., 1995; Hekengren & Hultmark, 1999; Lamberty et al., 2001a), or solely antifungal (Ijima et al., 1993; Fehlbaum et al., 1994; Schuhmann et al., 2003; Lamberty et al., 2001b). The antifungal activity of these AMPs has often been deduced from *in vitro* tests against opportunistic fungi (Levashina et al., 1995; Souhail et al., 2016; Fehlbaum et al., 1994; Yuan et al., 2007; Gao & Zhu, 2008; Tian et al., 2008, Zhang & Zhu 2009; Yang et al., 2006; Liu et al., 2016; Kim et al., 2001), but only a few times against ecologically relevant pathogens of insects (Hekengren & Hultmark, 1999; Tzou et al., 2002; Lu et al., 2016; Lamberty et al., 2001b). Moreover, the *in vitro* spectrum of activity of AMPs is not always transposable *in vivo* (Zanchi et al., 2017). This makes their contribution to insect fitness in case of a natural infection hard to predict, especially regarding peptides whose expression is not elicited by a fungal infection. In the mealworm beetle, *Tenebrio molitor* (Coleoptera: Tenebrionidae), it has recently been shown that an infection with *B. bassiana* activated the Toll pathway which lead to the expression of some antimicrobial peptides among such as the defensin Tenecin 1 and the coleoptericin Tenecin 2 (Yang et al., 2017), and that the knock down of this pathway decreased the survival of fungal infected beetles. However, the contribution of the constitutively expressed thaumatin Tenecin 3 (Makarova et al., 2016; Johnston et al., 2014; Yang et al., 2017) during *B. bassiana* infection in *T. molitor* is unknown.

It is commonly assumed that immune defense, including the production of antimicrobial peptides, comes at a cost (Johnston et al., 2014; Poulsen et al., 2002), and that its fitness benefits for the host have to outweigh these costs in order to evolve (Schmid-Hempel, 2005). Thus, we can expect the constitutive expression of Tenecin 3 to confer a benefit to *T. molitor* in the case of a natural fungal infection. The spectrum of activity of Tenecin 3 has been investigated *in vitro* towards several bacterial species whose growth were not affected, and against two opportunistic fungi, *Candida albicans* and *Saccharomyces cerevisiae* against which it was active (Kim et al., 1998; Kim et al., 2001).

In this study, we used the *Tenebrio molitor – Beauveria bassiana* model system to shed light on the contribution of a constitutively expressed antifungal peptide to host defense against a fungal infection. We used a gene knock-down approach by RNA interference on Tenecin 3, and monitored in parallel host fitness in terms of survival, and fungal fitness in terms of dynamics of within-host growth. We also performed an *in vitro* experiment in order to confirm the fungicidal effect of Tenecin 3 on *B. bassiana*.

Understanding the contribution of specific elements of the insect immune system in the resistance towards *B. bassiana* can help explain the success or failure of biocontrol strategies, as well as improve our understanding of the evolution of constitutive antifungal peptides as a defense mechanism.

## 2. Materials and Methods

### 2.1. Insect rearing

Larvae of the mealworm beetle (*Tenebrio molitor)* were purchased from a supplier and maintained at 25°C in the dark in plastic boxes (18 × 18 × 8 cm) at a density of 500 larvae in 400 g of wheat bran supplemented with rat chow. Fresh apple and albumin (from chicken egg white, Sigma) were added to the boxes every third day. Nymphs were collected every second day and kept apart until emergence. Newly emerged adult beetles were placed individually into grid boxes with wheat bran and a piece of filter paper. They received a piece of apple and ∼1g of albumin twice a week. Beetles of both sexes, aged 8-12 days post-hatching were used for the experiments.

### 2.2. Fungal cultures and conidia suspensions

A *Beauveria bassiana* strain (KVL 03-144) isolated from a natural infection of *Leptopterna dolobrata* (Homoptera: Miridae) from an agreoecosystem in Denmark (Meyling et al., 2009) was kept at −80°C in a culture collection at the University of Copenhagen before cultivation. Isolates were cultivated on quarter-strength Sabouraud Dextrose Agar + Yeast 10 % (SDAY) and incubated for 10 days at 23°C to allow for sporulation. Conidia were collected by scrapping the surface of the culture with a sterile loop and transferred in 1 ml PBS with 0.05% of Triton-X. The resulting solution was centrifuged twice at 23°C, 4000 rpm for 3.5 min and the supernatant discarded in order to remove agar and hyphae. The pellet was resuspended in 1 ml Phosphate Buffered Saline (PBS) + Triton-X 0.05%, and the conidia concentration assessed with a hemocytometer (Neubauer improved). The concentration of the inoculum was adjusted through serial dilutions before exposing the beetles. After each infection bout, the germination rate of the inoculum was assessed by plating 100 µL of a 10^5^ conidia/ml solution on quarter strength SDAY 10% plates. After incubation for 24 h at 23°C the germination of 3 × 100 conidia was counted. We discarded any batch of beetles that turned out to be infected with an inoculum of a germination rate lower than 90%.

### 2.3. Gene knock-down by RNA interference

We used RNA interference to knock down the expression of the *tenecin 3* gene. RNAi has been previously shown to be efficient in *Tribolium castaneum* (Coleoptera: Tenebrionidae) and *T. molitor* (Fabrick et al., 2009; Miller et al. 2012) and the effect to last at least 14 days (Zanchi et al., 2017). We generated double-stranded RNA (dsRNA) of a *tenecin 3* gene construct (Eurofins, Operon) which consisted in the sequence of the whole *tenencin 3* gene minus the sequence that was amplified by qPCR in order to confirm the efficiency of the knock-down of *tenecin 3* gene expression (Supplemental Information S1 figure S1). As a control, we used the dsRNA of the *Galleria mellonella* lysozyme, which has no homology of sequence with any known gene of *T. molitor* (Johnston & Rolff, 2015). The full sequences of the resulting products as well as the qPCR primers and products are given in the Supplemental Information S1. The template for dsRNA synthesis was amplified from the constructs by PCR (KAPA2G Fast ReadyMix, KAPA Biosystems) using gene-specific primers tailed with the T7 polymerase promoter sequence (Metabion International AG). After checking the length of our amplicon on a 2 % agarose gel and cleanup (PCR DNA Clean-Up Kit, Roboklon), the resulting amplicon was used as a template for RNA synthesis (High Yield MEGAscript T7 kit, Applied Biosystems/Ambion) according to the manufacturer’s recommendations. We then purified the RNA with a phenol-chloroform extraction and resuspended the pellet in a nuclease-free insect Ringer solution (128 mM NaCl, 18 mM CaCl2, 1.3 mM KCl, 2.3 mM NaHCO3). Before being used, the RNA was annealed by heating it up at 90°C and allowed to slowly cool down, in order to obtain dsRNA. We injected 500 ng of dsRNA per beetle at a concentration of 100 ng/µl in 5µL of insect Ringer solution. To do so, we chilled the insects on ice for 10-15 minutes before dsRNA injection, which was performed with a sterile pulled glass capillary inserted into the pleural membrane below the elytra, directly into the hemocoel. Care was taken that the needle was parallel to the anterior/posterior axis of the beetle to avoid injury of the organs. Beetles that received the Tenecin 3 dsRNA will be referred to as “Ten3KD” whereas beetles that received *G. mellonella* dsRNA will be referred to as “control”.

### 2.4. Fungal exposure and maintenance of infected beetles

Exposures to *B. bassiana* were performed by applying a 0.5 µl droplet of conidia/PBS-TritonX solution prepared as described above on the intersegmental membrane between the sclerotized parts of the sternum and the abdomen. We performed the exposures 7 days after the injection of dsRNA. Care was taken so that the whole droplet was adsorbed on the membrane before placing the beetles individually at 23°C in medicine cups (Carl Roth GmbH) containing a 2×2cm wet filter paper in order to keep the humidity level high in the cup and stimulate *B. bassiana* conidia to germinate. Beetles were provided with food (ca. 5 g sterilized rolled oats) a day later in order to avoid the transfer of conidia in the medium. The filter papers were replaced every day, and the medicine cups along with the food were replaced every four days.

### 2.5. Mortality bioassays

We tested the effect of three different concentrations of *B. bassiana* conidia on beetles survival: 1×10^5^, 5×10^5^ and 1×10^6^/ml, corresponding to 50, 250, 500 conidia applied per beetle, since previous laboratory experiments showed that these concentrations span the LC50 (3×10^5^ conidia/ml) of the strain KVL 03-144 of *B. bassiana* (S. Maistrou, unpublished). As procedural control we included a group of beetles that each received only a 0.5 µl PBS-0.05% Triton-X solution. The beetles were maintained as described above and their mortality was checked daily for 14 days after exposure. All dead beetles exposed to *B. bassiana* developed mycosis after death. Each treatment group consisted of 20 beetles and the bioassay was repeated twice. The final sample sizes are presented in the Supplemental Information S2 Figure S2.

### 2.6. Recovery of hyphal bodies from the beetles hemolymph

We chose to perform this experiment on beetles exposed to 0.5 µl of 5×10^5^ conidia/ml since this was the concentration which yielded the biggest effect on mortality between Ten3KD and control beetles (Figure 1). Beetles exposed to *B. bassiana* were split into two groups of ∼100 beetles/treatment (control or Ten3KD). Since hemolymph collection was an invasive process, it was performed only once per beetle. Therefore, at each time point (1, 2, 3, 4 and 5 days post infection) we collected the hemolymph of ∼20 randomly selected beetles of each of the two treatment groups.

**Figure 1:**
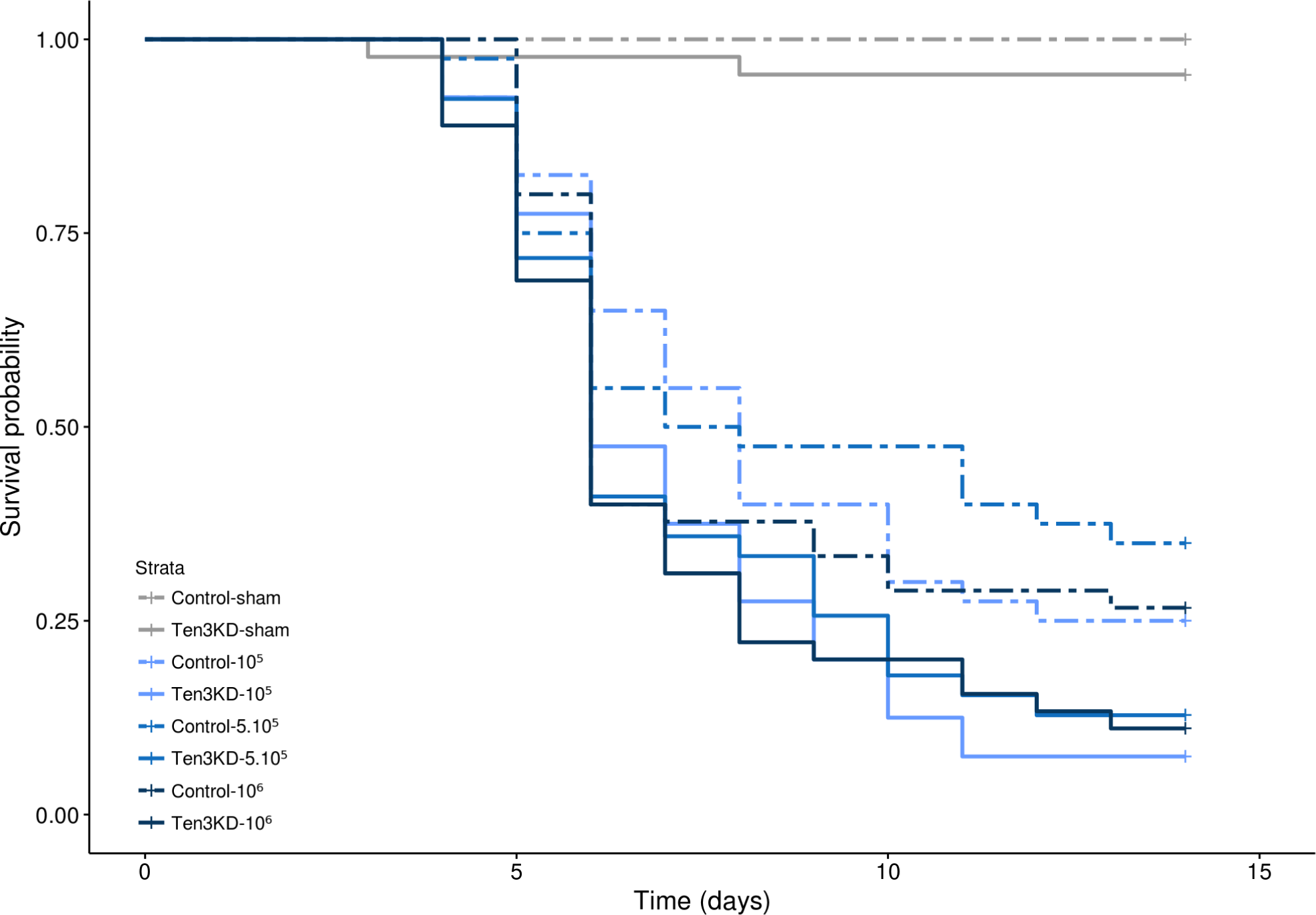
Tenecin 3 improves the survival of *Tenebrio molitor* to *Beauveria bassiana*. Kaplan-Meier curve showing the survival over 14 days of the control (dashed lines) and Tenecin 3 knock-down (solid lines) beetles either sham (in grey) or exposed to *Beauveria bassiana* (in blue, lighter to darker shades indicating lower to higher concentrations : light blue = 10^5^ conidia/mL; medium blue = 5.10^5^ conidia/mL; dark blue = 10^6^ conidia/mL). Only the knock-down treatment had a significant effect on survival of the beetles (dashed vs. solid lines, Х²_2,247_ = 8.13; p = 0.017).

Before collection, beetles were chilled on ice (10-15 minutes) in a glass tube containing a piece of cotton soaked in ethyl-acetate and briefly dipped in 70% ethanol to remove any contamination from the cuticle. An incision was made above their genitals before inserting a hypodermic needle of a syringe containing 0.5 ml PBS which was subsequently perfused. The resulting haemolymph/PBS extract was collected into a microcentrifuge tube kept on ice. 100 μL of this suspension was plated on SDA + 5 mg/ml tetracyclin using sterile glass beads, and incubated at 23°C for 36 h (we established beforehand that tetracycline did not affect the growth of blastospores in terms of speed of germination and density, data not shown). The number of colony forming units (CFU) on each plate was counted and used as a proxy to assess the concentration of *B. bassiana* hyphal bodies in the hemolymph of each individual beetle at a given time point. Few beetles died before the end of the experiment, which was compensated in our sample size by the fact that we injected slightly more than 100 beetles at the start. Some cadavers developed mycosis as expected (i.e. succumbed to *B. bassiana* infection); there were 1 beetle in both the control and the Ten3KD while 2 beetles in the control and 3 in the Ten3 KD treatment died without showing mycosis symptoms. The final sample sizes are presented in the Supplemental Information S2 Figure S3.

### 2.7. *In vitro* effect of Tenecin 3 on *Beauveria bassiana*

#### 2.7.1. Growth and collection of blastospores

500 μl of conidial suspension (10⁷ conidia/ml) was added into a flask containing 100 ml of Sabouraud Dextose broth + 10% yeast extract. The solution was incubated for 72-96 h at 23 °C with agitation at 150 rpm, in order for the conidia to germinate. After incubation, the solution was filtered through a sterile filter paper (595 grade, Schleicher & Schuell) in order to recover the blastospores and discard the mycelium. The resulting solution was washed by centrifugation three times at 10 000 g and 25 °C for 10 minutes, and the pellet resuspended in 1ml PBS. The concentration of blastospores in this solution was assessed in a hemocytometer, and adjusted to 5.10^6^ blastospores/ml.

#### 2.7.2. Survival of blastospores *in vitro* against a recombinant Tenecin 3

We first established a growth curve of the *B. bassiana* strain KVL 03-144 in our experimental conditions in order to focus on a time point which was in the exponential phase for the rest of the experiments. The protocol and the growth curve are presented in the Supplemental Information S3.

We chose to carry out the rest of the next experiment on the 6 hour time point, corresponding to the beginning of the exponential growth phase. We inoculated 19.5 µl of Sabouraud Dextrose broth with 0.5 μl of blastospore solution (10^3^ blastospores) in the wells of a 96 well plate (Sarstedt). We added 5µl of a solution of water and recombinant Tenecin 3 resulting in a final concentration of 0.05, 0.1, 0.2, and 0.4 ng/μl of Tenecin 3. After incubation at 25 °C and agitation at 220 rpm for 6 h, we suspended the content of the wells in PBS up to 100 µl and plated 100 µl of serial dilutions by a factor of 10 and 100 on SDA. We incubated the Petri dishes at 23°C in the dark for 48 hours after which we counted the number of CFU. As a control, we replaced Tenecin 3 with bovine Serum Albumin (BSA) at the same concentrations. The process was repeated 5 times, with conidia originating from different culture plates.

### 2.8. Data analysis

All statistical analyses were performed using the R software (R Core Team, 2016). The survival of the beetles after *B. bassiana* exposure was analyzed with the “survreg” function of the ‘survival’ package (Therneau, 1999), since the hazards were not proportional between treatment (checked with the “coxzph” function). We checked whether survival of the beetles was affected by the concentration in conidia of *B. bassiana* used for exposure and their knock-down treatment, as well as the interaction between them and the replicate as random factor, and with an exponential distribution. Both sham infected treatments showed a very low mortality and impaired the fit of the model, therefore we decided to exclude them during the analyses, but show them on the figure.

The concentration of hyphal bodies present in the hemolymph of the beetles was analyzed with a Generalized Linear Model fitted with a negative binomial distribution with the package ‘MASS’ (Venables & Ripley, 2002). We tested whether the concentration of hyphal bodies was explained by the knock down treatment and the time as well as their interaction, in order to highlight the dynamics of the progression of the infection.

The *in vitro* growth of blastospores in the presence of Tenecin 3, BSA or alone was compared using a Generalized Linear Mixed Model with a Poisson distribution corrected for overdispersion with the package ‘lme4’ (Bates, 2015), including the treatment (protein added, either BSA or Tenecin 3) as well as the protein concentration as explanatory variables as well as the interactions between them, and the replicate as a random factor.

For all the analyses, we chose which distribution to use by checking the distribution of the residuals and comparing the estimates of the models with our data with the package ‘visreg’ (Breheny & Burchett, 2017). We then selected the best model by comparing the Akaike’s Information Criterion (AIC) of the full models including interactions to all the nested models and the null model. We kept as the best models the ones with the lowest AICs (Akaike, 1976). Post hoc comparisons when relevant can be performed by comparing the 95% confidence intervals around the estimates of the models on the figures provided in the Supplemental Information S4. A difference between two treatments is deemed significant when the confidence intervals do not overlap on more than half of their length, as advised by Cumming (2009). The figures in the main body of the manuscript were made using the ggplot2 package (Wickham, 2009), and the figures of the Supplemental Information were obtained with the package ‘effects’ (Fox & Hong, 2009).

## 3. Results

### 3.1. Survival of the beetles after *B. bassiana* exposure

There was no effect of the concentration of conidia applied on the beetles for infection either in interaction with the treatment (concentration*treatment: Х²_6,243_ = 9.12; p = 0.17) or alone (Х²_3,246_ = 0.85; p = 0.84) across the range of concentrations we checked. The knock-down of Tenecin 3 significantly reduced survival of the beetles after *B. bassiana* exposure (Х²_2,247_ = 8.13; p = 0.017; LT50 control = 7.5 days; LT50 Ten3KD = 5.5 days).

We then decided to keep investigating the effect of the knock-down treatment in beetles exposed to 0.5 μl of 5×10^5^ conidial/ml (∼ 250 conidia) on the *in vivo* progression of the infection over 5 days, after which the mortality of the beetles would have biased our sampling between treatments.

### 3.2. Progression of the infection in the beetle host

Very few hyphal bodies could be recovered from the early time points (less than 10 and less than 50 on the first and second day post infection respectively, see Figure 2). Both control and Ten3KD treatments showed similar dynamics of the progression of the infection, i.e. there was no interaction between time and treatment on the number of hyphal bodies recovered from the beetles (time*treatment : Х²_3,202_ = 1.28; p = 0.257). However, both the treatment and time affected the concentration of hyphal bodies present in the beetles as simple effects. Ten3KD beetles contained overall more hyphal bodies than control beetles (treatment : Х²_1,203_ = 7.61; p = 0.00578). As expected, the concentration of hyphal bodies increased over time (time : Х²_1,203_ = 211.205; p < 0.0001, see Supplemental Information S4 Figure S5 for post hoc comparisons).

**Figure 2:**
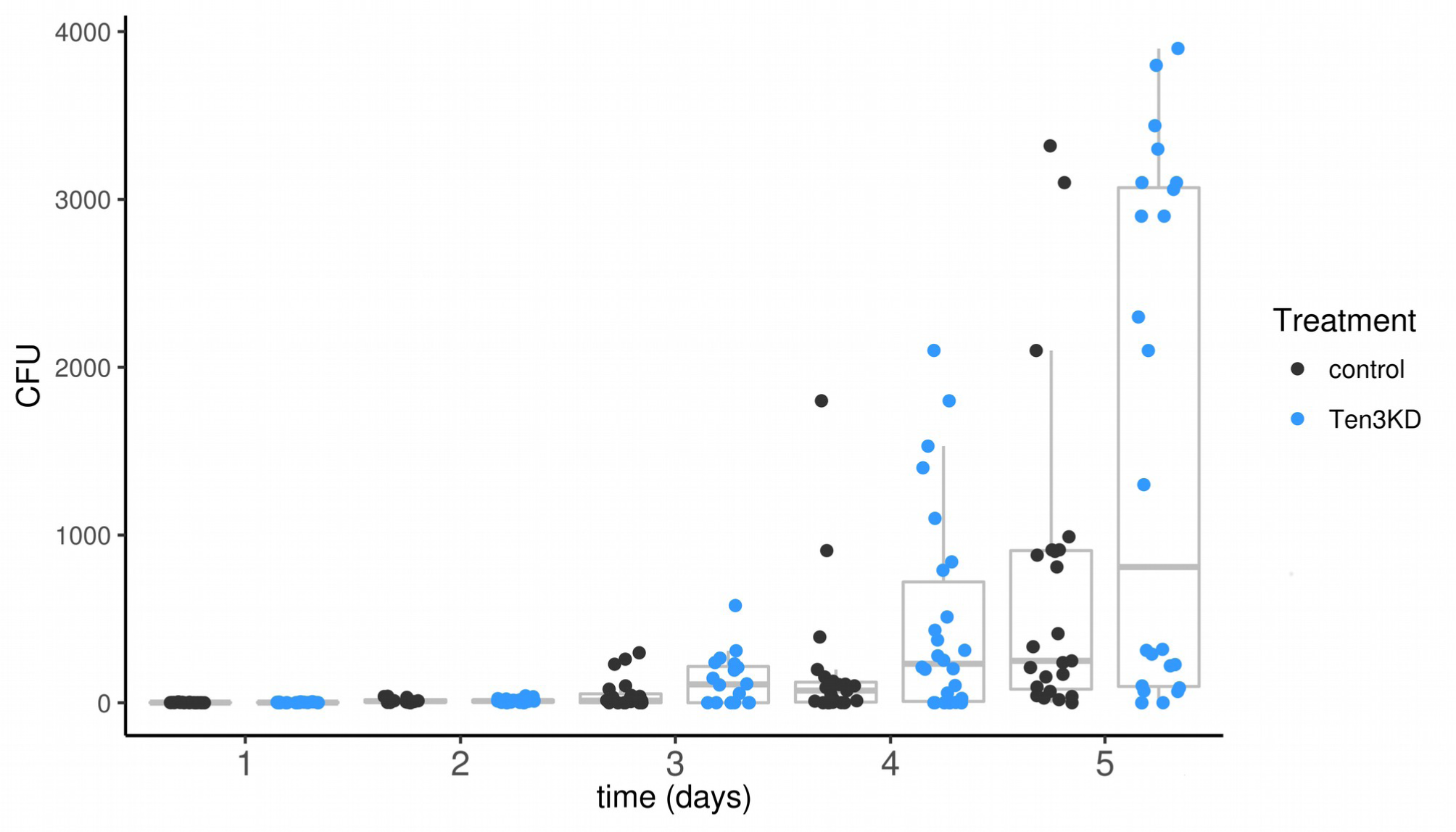
Tenecin 3 decreases the fungal load of *Beauveria bassiana* in the hemolymph of *Tenebrio molitor*. Box-dotplot representing the number of CFU of *B. bassiana* recovered from the hemolymph of infected beetles according to the time after infection. Each dot represents a data point of control (grey) and Tenecin 3 knock-down (in blue) beetles. The boxplots represent the first to the third quartiles around the median (horizontal grey line), and the vertical bars the 1.5 interquartile of the lower and upper quartiles. Both time and treatment affect the fungal load in the hemolymph of the beetles, see Supplemental Information S4 Figure S5 for post-hoc comparisons.

Moreover, an interesting pattern emerges from the distribution of our data. The quantity of hyphal bodies present in the beetles seems to evolve over time towards a bimodal distribution (Figure 2), which is very obvious at 5 days post infection. By setting a cut-off at 600 CFU at this time point we can determine that in the control, 39.1% of the data points are above this threshold, versus 50.0% for the Ten3KD. By looking at the survival curves of the two treatments in the corresponding concentration (5.10^5^ conidia/mL, Figure 1 in medium blue) it appears that they diverge at 6 days post-infection and remain relatively parallel after this time point. The proportions of dead beetles at this point are 45.0 and 59.0% for control and Ten3KD respectively (Figure 1). While the 24 h time frame of our sampling does not allow us to capture the concentration of hyphal bodies present in the hemolymph of the beetles immediately prior to death, we can observe that these values are not too different from the proportion of the beetles who carried more than 600 CFU at day 5 (Figure 2).

### 3.3. In vitro killing of *B. bassiana* blastospores by recombinant Tenecin 3

We investigated whether the lower concentration in hyphal bodies in the hemolymph of control beetles compared to Ten3KD beetles could be due to a direct effect of Tenecin 3 on *B. bassiana* hyphal bodies. We exposed blastospores (hyphal bodies generated *in vitro*) to a recombinant Tenecin 3 or to BSA as a control at different concentrations for 6 hours, and compared the concentration of blastospores recovered after incubation.

Tenecin 3 affected the number of blastospores in the medium differently than BSA at higher concentrations (0.2 and 0.4 ng/µl : concentration * treatment : Х²_8,34_ = 552.56; p < 0.001, see Supplemental Information 4 Figure S6 for post-hoc comparisons). The number of blastospores was lower when incubated at both 0.2 and 0.4 ng/µl of Tenecin 3 compared to the inoculum, suggesting a fungicidal effect of Tenecin 3 on *B. bassiana* (**Figure 3**).

**Figure 3:**
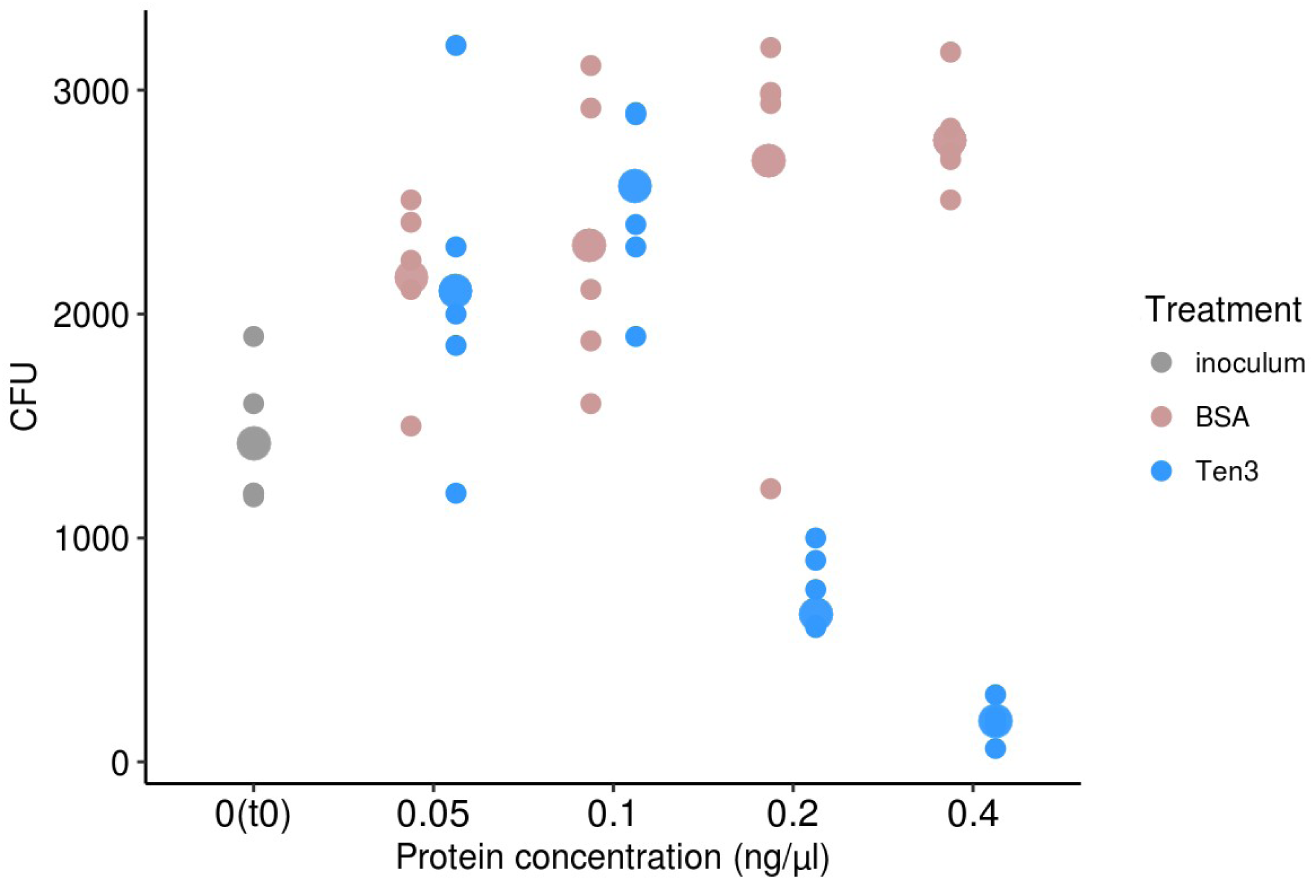
Recombinant Tenecin 3 has an antifungal activity against the blastospores of *Beauveria bassiana in vitro*. Dotplot representing the number of CFU retrieved after *in vitro* incubation of blastospores for 6 hours with either Bovine Serum Albumin (BSA, in pink) or recombinant Tenecin 3 (in blue). Five replicates were performed, each dot represents a data point and the thicker dot represents the mean. The blastospore solution (around 10^3^ conidia) was plated after inoculation and is represented in grey as “0(t0)”, meaning a concentration of 0 and time point 0. Starting from 0.2ng/µL, Tenecin 3 has a fungicidal effect on the blastospores of *B. bassiana*, see Supplemental Information S4 Figure S6 for post hoc comparisons.

## 4. Discussion

We found that the constitutively expressed antifungal peptide Tenecin 3 increases beetle survival to a *B. bassiana* infection. This is mirrored by a clear antifungal effect of Tenecin 3 on *B. bassiana* blastospores *in vitro*.

Antimicrobial peptides of insects are well studied in the context of bacterial infections, in which case they are mostly inducible (Bulet & Stöcklin, 2005). By comparison, *in vivo* studies on antifungal peptides in the context of fungal infections established through the natural route are scarce. The constitutive expression of Drosomycin, an otherwise inducible antifungal peptide, in adult *Drosophila melanogaster* (Diptera: Drosophilidae) mutants deficient for both the Toll and Imd pathways, restored the wild-type survival to opportunistic or human pathogenic fungi injected in the flies but did not improve survival to a natural infection with *B. bassiana* (Tzou et al., 2002). Termicin, an antifungal peptide constitutively expressed in the salivary glands of some species of termites has been shown to increase the survival of *Reticulitermes flavipes* to a *Metarhizium anisopliae* infection (Hamilton & Bulmer, 2012). The first result can seem counterintuitive considering that Drosomycin is elicited following *B. bassiana* infection (Lemaître et al. 1997). In the second case, it is likely that the selection pressure exerted by fungal pathogens is constant in the environment of these species of termites, making the evolution of an external constitutive antifungal defense relevant (Bulmer & Crozier, 2004; Adler & Karban, 1994). In *T. molitor*, by performing knock-downs of the Toll pathway, Yang et al. (2017) highlighted the importance of inducible immune defenses in the survival to *B. bassiana*, although the authors did not directly knock-down the inducible AMPs suspected to be involved, whereas the role of the constitutive Tenecin 3 was not investigated. Even though the pressure exerted by *B. bassiana* on *T. molitor* in its natural habitat is unknown, *B. bassiana* is widespread in most ecosystems investigated (e.g. Meyling et al. 2011; Hajek & Meyling 2018; Eilenberg et al. 2015) and we could expect that similarly to what is observed in the aforementioned termite species, the fact that a constitutively expressed antifungal peptide improves the survival of *T. molitor* during a *B. bassiana* infection indicates that Tenecin 3 might have evolved as a way to fight off fungal a constant selection pressure exerted by fungal pathogens in the environment. If this is the case, the changes of its expression patterns during development (Lee et al., 1996) might even indicate that this pressure is stage-dependent. Since other AMPs are induced in *T. molitor* following *B. bassiana* infection through the Toll pathway (Yang et al., 2017), a future lead to follow could be to perform knock-down on these peptides simultaneously, which would shed light on the interplay of constitutive and inducible immune defenses.

Our study also adds support to previous observations made in *D. melanogaster* that the evolution of the pathogen load inside the host over time can show a bimodal outcome, where one category of hosts carry a low level infection while the other category will die of high infection levels (Clemmons et al., 2015; see Duneau et al., 2017 for data & review). Similarly to Duneau et al. (2017), the quantity of hyphal bodies present in the beetles at a certain time point seems to be responsible for their mortality, and suggests that there is a threshold pathogenic load which needs to be reached by the fungus in order to kill the host. However the overall pattern of progression of the infection differs compared to these previous studies : while after injection of bacteria in both *D. melanogaster* and *T. molitor* there is an immediate dramatic decrease of the pathogen load in the host, our present infections seems more to build-up from a low load in the hosts hemolymph. In these previous studies a high concentration of bacteria was directly injected into the hemolymph, bypassing the cuticular barrier. With this protocol, Zanchi et al. (2017) did not find an effect of the knock down of various AMPs on the early stages of the infection, but instead an increased bacterial load and dispersion of the bacterial counts in later time points after infection. Our present study therefore confirms this result in the case of a natural infection, which can be considered like an ecologically relevant setting.

To conclude, we show that the direct antifungal activity of Tenecin 3 on *B. bassiana* protects *T. molitor* against the internal progression of infection by this fungus. While many current studies focus on insect defenses which prevent the infection from being initiated, the role of the immune system of the hemolymph in host resistance to entomopathogenic fungi should not be underestimated. Understanding the contribution of specific elements of the insect immune system in the resistance towards *B. bassiana* can help explain the success or failure of biocontrol strategies, as well as improve our understanding of the evolution of constitutive antifungal peptides as a defense mechanism.

## Acknowledgements

We thank P. R. Johnston, O. Makarova & A. Rodriguez Rojas for useful insight and comments on our experiments, and N. Demandt for useful advice on statistical analyses.

## Funding

This research was funded by the ERC (Grant n° 260986) and the University of Copenhagen.

