## Supplementary Materials for "A constitutively expressed antifungal peptide protects *Tenebrio molitor* during a natural infection by the entomopathogenic fungus *Beauveria bassiana*"

### Supplemental Information S1 : gene knock-down by RNA interference

#### Sequence of the *tenecin 3* gene construct.

The synthetic gene construct used in this study is based on the sequence of the *tenecin 3* gene obtained from Jung et al. (1995) from which we omitted the sequence used to follow the expression of the *tenecin 3* gene by qPCR. The product of its amplification was used as template for the synthesis of dsRNA.
The sites of attachment of the qPCR primers are highlighted in pink (product of 77bp), the ones of the attachment of the primers used for the amplification of the construct before transcription in blue (254 bp).

NCBI accession number U21482

CCAAAATGAAAACATTCGTGATTTGCTTGATTCTGGTGGTCGCCGTTTCGGCAGC
TCCTGACCATCACGACGGACATCTGGGTGGTCACCAAACCGGTCACCAAGGCGGC
CAACAGGGTGGTCATCTGGGGGGTCAACAGGGTGGACACCTAGGGGGTCACCAG
GGCGGCCAACCAGGCGGACATTTAGGAGGCCATCAGGGCGGAATCGGAGGCACC
GGAGGGCAGCAACACGGGCAGCATGGACCTGGGACCGGTGCAGGACACCAAGGA
GGGTACAAGACACATGGTCATTAATGTGAATTATATACATATACAGAGTGGCCAT
AAAATTGTCA (337 bp)

The primers used for the amplification of the construct were tailed with the T7 polymerase promoter sequence (in bold) :

forward : **TAATACGACTCACTATAGGGAGA**CCAAAATGAAAACATTCGTGATT

reverse : **TAATACGACTCACTATAGGGAGA**GGTCCCAGGTCCATGCTG

Which yielded a final product of 300 bp.

We used Galleria mellonella lysozyme, used as a procedural control for dsRNA injection, SwissProt accession number P82174

### Verification of RNAi knock-down efficiency

The CT values of the tenecin 3 gene and the reference gene (RPL27A) were assessed over 40 PCR cycles (StepOneTM RealTime PCR System, Applied Biosystems) with the following parameters:
Holding stage 95°C, 20 seconds
Denaturation 95°C, 1 second
Extension 60°C, 20 seconds

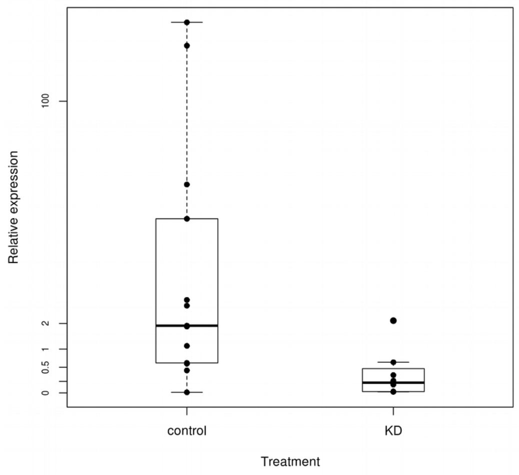


**Figure S1 :** Relative expression of the *tenecin 3* gene compared to an internal control (ribosomal protein RPL27A gene) in control and *tenecin 3* knockdown (KD) beetles (n = 10 per treatment, from various time points following exposure, 24h, 3 and 4 days). There is a significant average 115 times reduction of expression of the *tenecin 3* gene in KD individuals compared to control (negative binomial GLM : 0.43 vs. 49.73 ; $F_{1,19}$ = 14.88 ; p = $1.14e^{-4}$)).
