## Supplementary Materials for "A constitutively expressed antifungal peptide protects *Tenebrio molitor* during a natural infection by the entomopathogenic fungus *Beauveria bassiana*"

### Supplemental Information S2 : Experimental designs

#### Figure S2 : Mortality bioassays

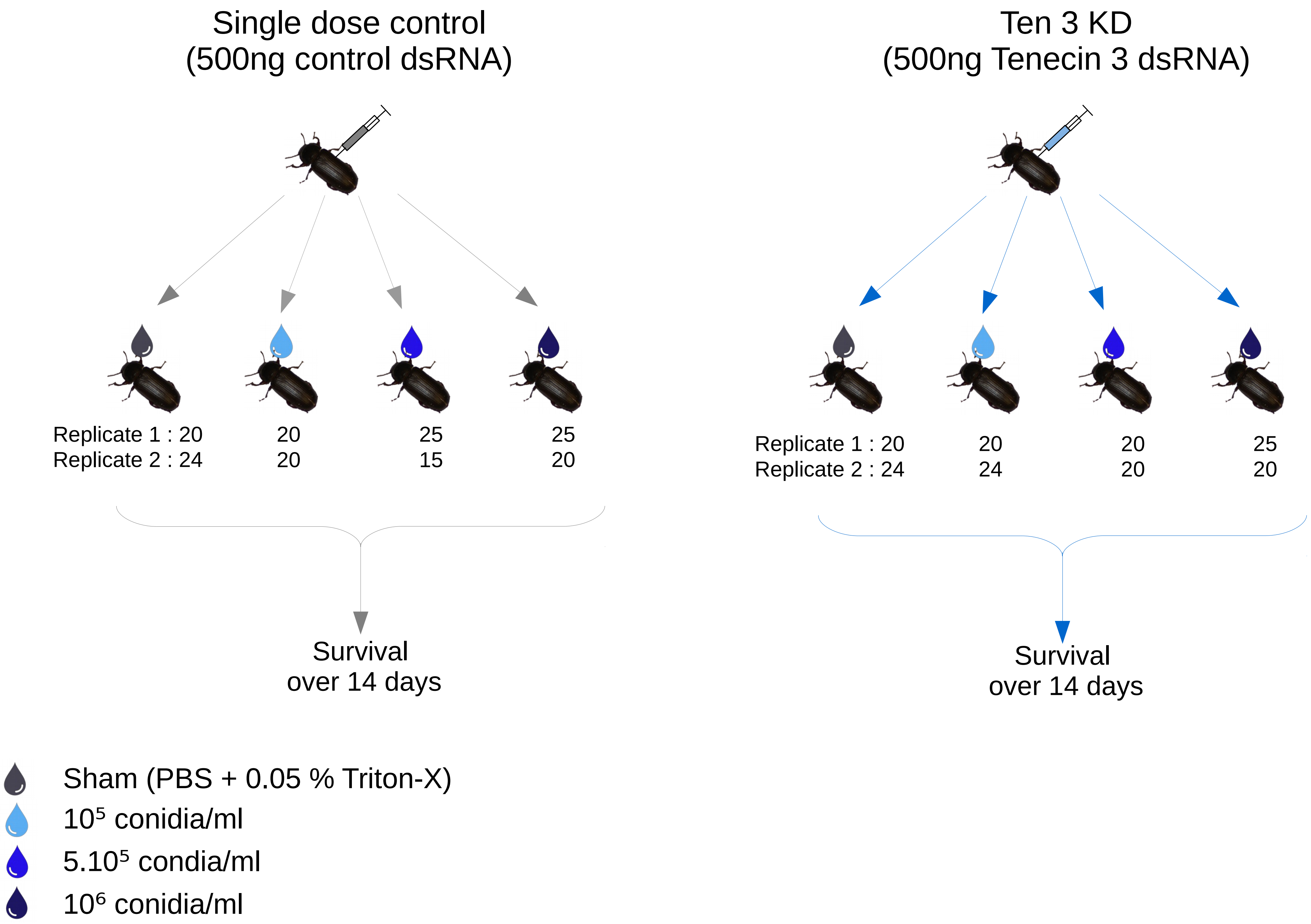

### Figure S3 : Recovery of hyphal bodies from the beetles hemolymph

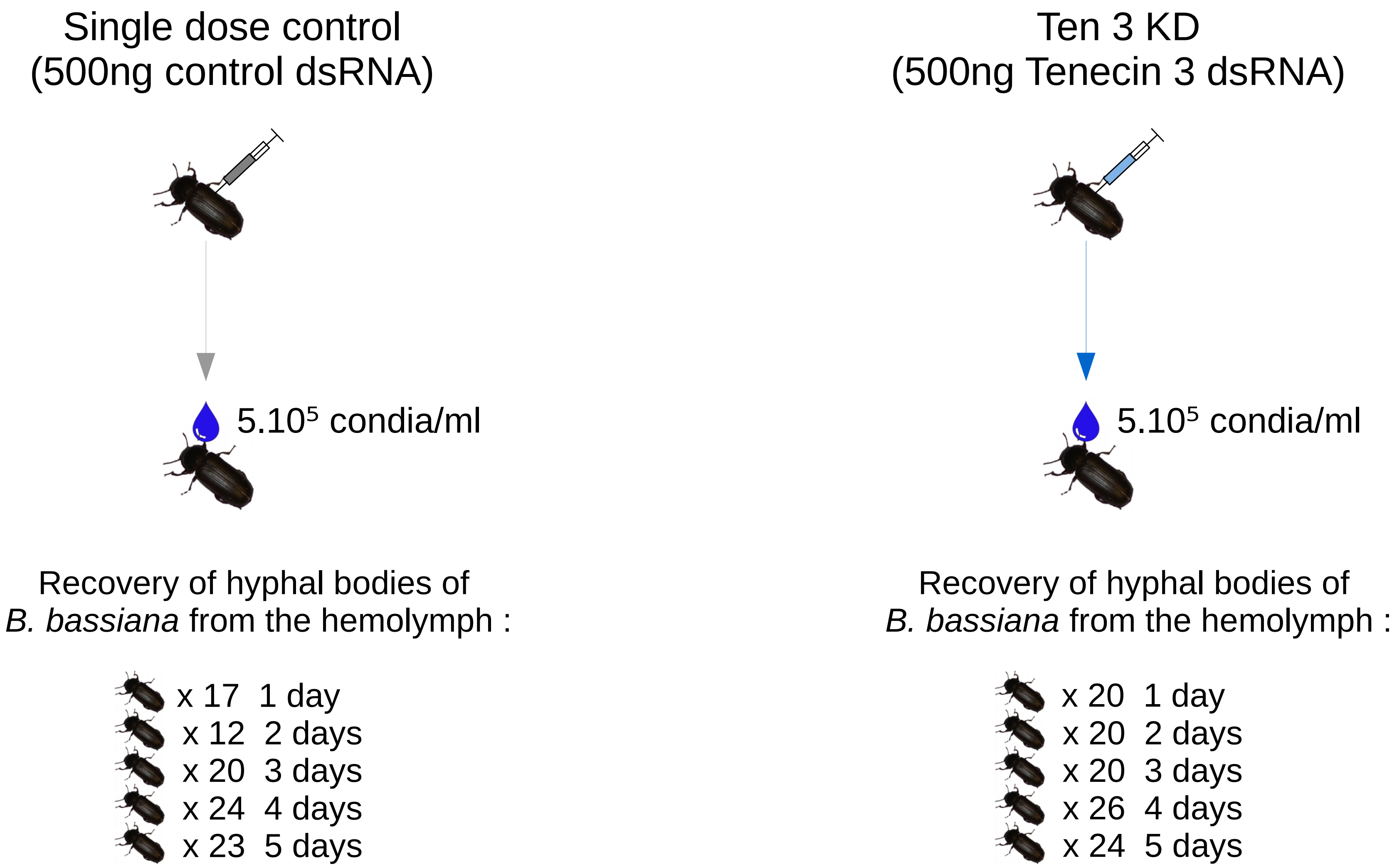
