## Supplementary Materials for "A constitutively expressed antifungal peptide protects *Tenebrio molitor* during a natural infection by the entomopathogenic fungus *Beauveria bassiana*"

### Supplemental Information S3 : growth curve of *Beauveria bassiana* KVL 03-144 blastospores *in vitro*

A previous paper by Kim et al. (2001) showed that Tenecin 3 activity against *Candida albicans* relied on its internalisation in growing cells by an active process. We therefore determined what time frame corresponded to the exponential phase of growth of the blastospores of our strain in our experimental conditions. To do so, we added 0.5 µl of blastospores suspension at $5.10^6$ blastospores/ml (~ 1000 blastospores) in 24.5 µl of Sabouraud Dextrose broth + 10% yeast extract in a 96 wells plate. We incubated the plate at 25 °C and recovered its content after 3, 6, 9, 12 or 24 hours after inoculation. We adjusted the volume of the incubation product up to 100 µl with PBS, plated the content of the wells on Sabouraud Dextrose Agar + 10% yeast extract, and incubated the Petri dishes at 23°C for 48 hours in the dark, after what we counted the number of Colony Forming units (CFU) present on the plates. The process was repeated three times, and an incubation-time of six hours was chosen for the experiment in section 3.3.

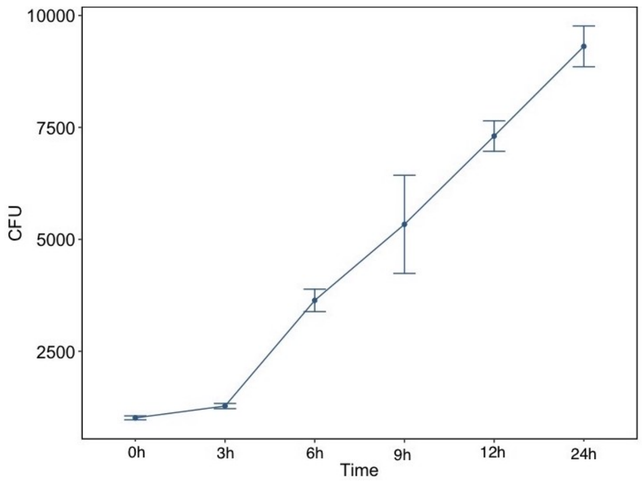


**Figure S4 :** Growth curve of *B. bassiana* KVL 03-144 in Sabouraud Dextrose broth + 10% yeast extract. The dots represent the mean number of CFU of *B. bassiana* recovered from 3 replicates, the vertical bars represent the standard deviation for each time-point.
