## Supplementary Materials for "A constitutively expressed antifungal peptide protects *Tenebrio molitor* during a natural infection by the entomopathogenic fungus *Beauveria bassiana*"

### Supplemental Information S4 : details on the statistics

**Figure S5 for post-hoc comparisons in the section 3.2. :**Effect plot of the GLM on the number of CFU of *Beauveria bassiana* hyphal bodies recovered from the hemolymph of *Tenebrio molitor* according to a) treatment and b) time, both these factors having a simple effect on the number of CFU recovered. The dots represent the estimates of the GLM, the vertical bars the 95CI. Post-hoc comparisons can be performed by looking at the overlap of the confidence intervals of the different treatments.

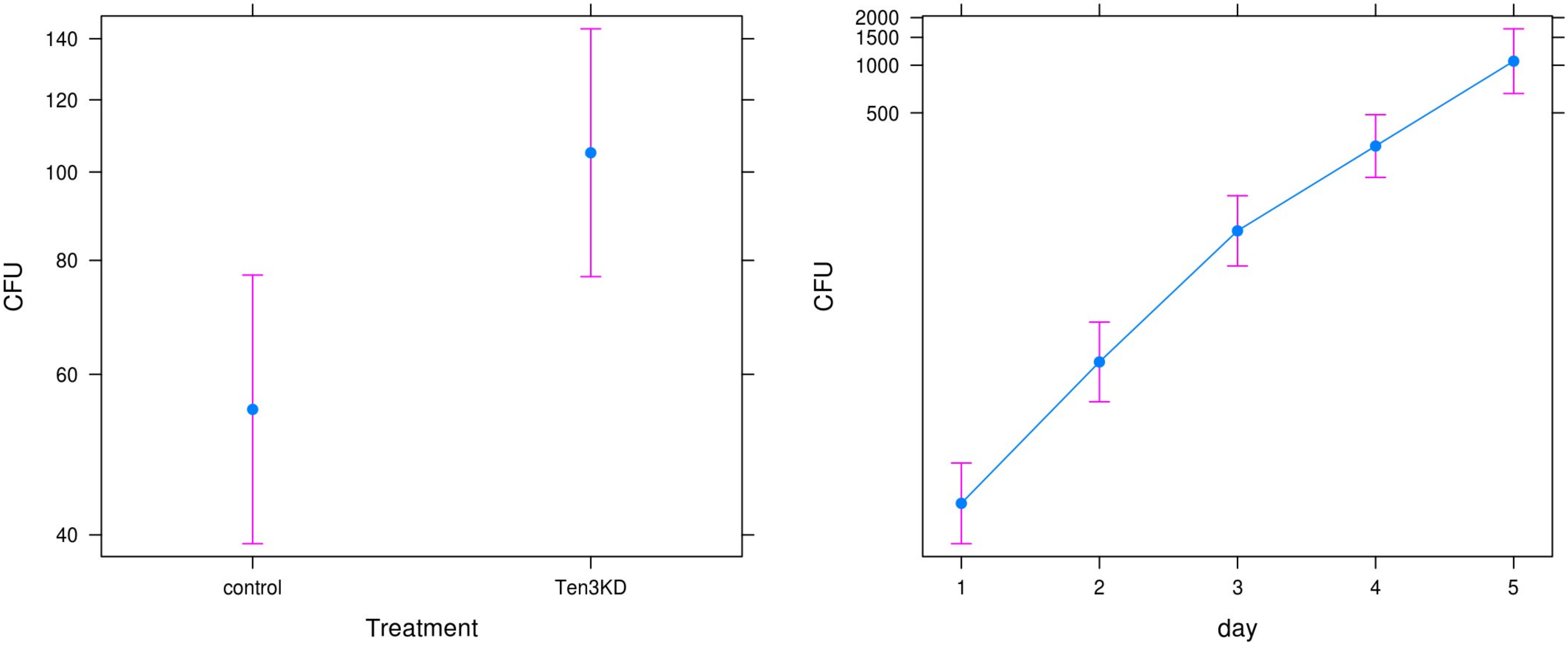

**Figure S6 for post-hoc comparisons in the section 3.3. :** Effect plot of the GLM on the number of CFU of *B. bassiana* blastospores recovered from the inoculum (0(t0)) and from the growth medium 6 hours after incubation with either BSA or recombinant Tenecin 3 at different concentrations. The dots represent the estimates of the GLM, the vertical bars the 95CI. Post-hoc comparisons can be performed by looking at the overlap of the confidence intervals of the different treatments.

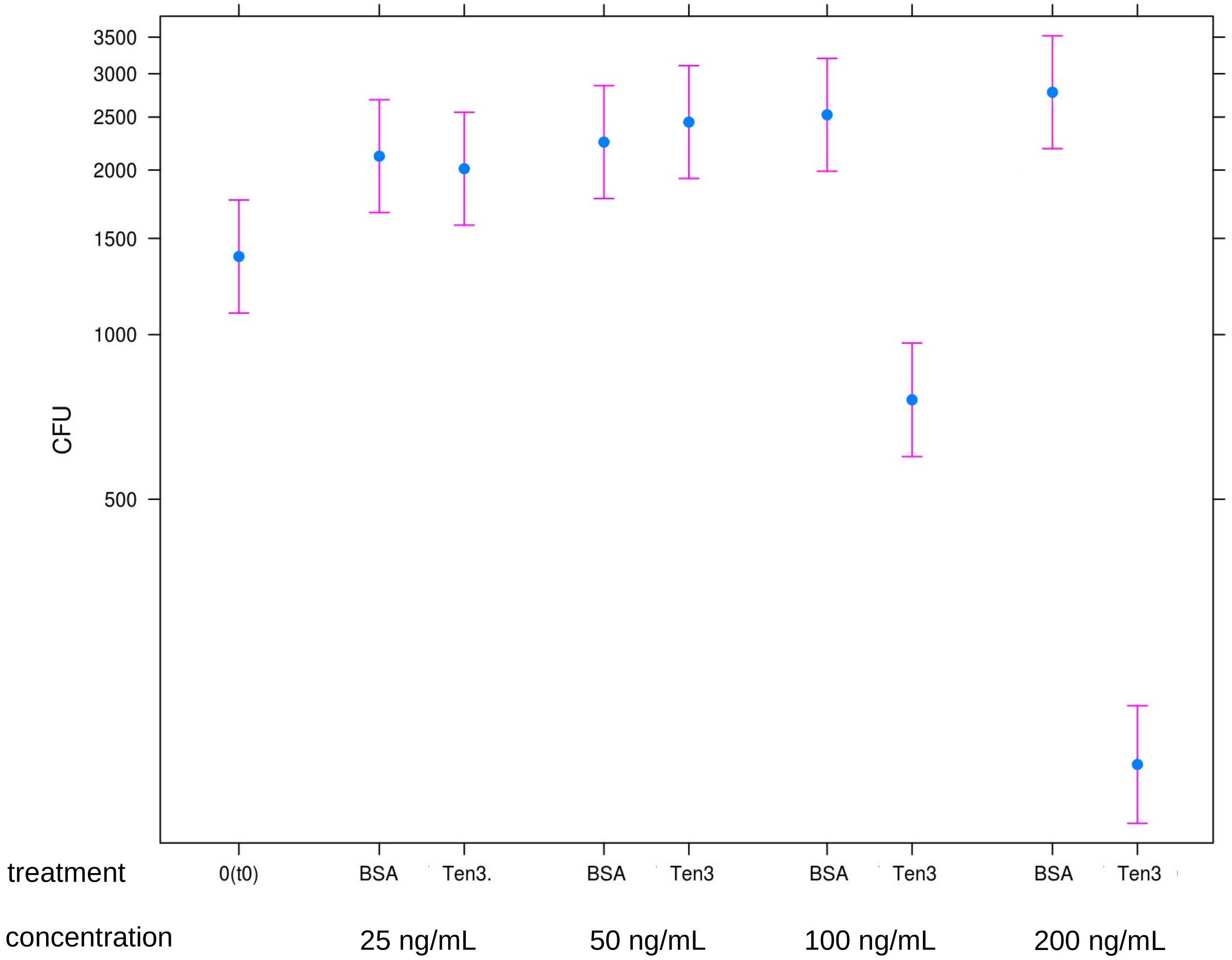
